# Efficiently determining membrane-bound conformations of peripheral membrane proteins using replica exchange with hybrid tempering

**DOI:** 10.1101/2024.05.04.592548

**Authors:** Chandramouli Natarajan, Anand Srivastava

## Abstract

Accurately sampling the membrane-bound conformations of a peripheral membrane proteins (PMP) using classical all-atom molecular dynamics simulations (AAMD) is a formidable enterprise due to the rugged free energy landscape of the protein-membrane system. In general, AAMD-based extraction of binding geometry requires simulations of multiple systems with different initial user-defined binding poses that may not be exhaustive. As an alternative, advanced sampling methods are also applied to elucidate the membrane-binding mechanism of PMPs but these techniques are generally computationally expensive and often depend on choice of the collective variables (CV). In this work, we showcase the utility of CV-free replica exchange with hybrid tempering (REHT) method in capturing the membrane-bound conformations of PMPs by testing it on the Osh4 amphipathic lipid-packing sensor (ALPS) motif, a 27 amino-acid membrane binding peptide. We show that REHT samples all the membrane-bound conformations of the Osh4 ALPS peptide at their correct populations and does it in a highly efficient manner with minimum computational time. We clearly show that, out of the two significant conformations, the peptide prefers horizontal conformations over vertical ones. In both the conformations, REHT captures all the vital residue-wise membrane contacts. The transition between the two configuration is not uncommon as our calculations reveal a ∼ 2 kT free energy difference between the two conformations. Interestingly, from our simulations we also find that the transition from vertical to horizontal conformation involves limited unfolding the main helix’s last turn. From our findings, we conclude that REHT samples the membrane-bound conformations of Osh4 ALPS peptide very efficiently and also provides additional insights and information that are often not available with regular piece-wise AAMD simulations. The method can be used as an efficient tool to explore the membrane-binding mechanisms of PMPs.

## 1 Introduction

Peripheral membrane proteins (PMP) are a class of membrane proteins that localize at the surface of the membrane. The function of PMP depends both on its membrane affinity and as well as the membrane-bound conformations. Also, the proteins carry different membrane interacting motifs and domains that allow them to interact with specific membranes to perform their function.^1–3^ Therefore, accurate characterization of PMP membrane binding mechanisms is crucial to understanding their function. Experimental methods such as cryo-EM, solid state NMR and EPR are applied, when necessary, to determine the membrane-bound structure of the PMP but often due to the membrane and other experimental constraints, the membrane-bound conformation and dynamics information are low-resolution in nature.^4,5^ Along with experimental methods, computational algorithms such as all-atom molecular dynamics (AAMD) and related simulations have been effectively used to supplement the experimental information and elucidate the residue-level membrane association information of PMP.^6–10^

However, expounding the equilibrium properties of the membrane-binding characteristics of a PMP using simulations is non-trivial.^11–13^ Generally, a protein-membrane system exhibits a rugged free energy landscape with multiple minima separated by high free energy barriers. Hence, classical (brute-force) AAMD simulations may not be adequate to sample the entire conformational landscape of the membrane binding process of a PMP. Such a shortcoming could be overcome by using an enhanced sampling technique like replica exchange simulations (REX).^14,15^ A few studies have shown that replica exchange with solute tempering (REST) explores the protein-membrane interactions faster and better than classical MD or traditional replica exchange molecular dynamics (REMD).^16–19^ Recently, we developed a variant of REX called replica exchange with hybrid tempering (REHT) for the exhaustive sampling of intrinsically disordered proteins (IDPs).^20^ In this method, the differential and optimal heating of both the protein (solute) and water (solvent) helps the protein cross significant free energy and entropic barriers. Hence, the IDPs could escape metastable states quickly and explore all conformational basins. Similarly, the process of membrane binding by proteins may be strewn with metastable states and considerable free energy barriers. Such traps might capture the membrane-bound protein in one state and prevent the exploration of other membrane-bound states. ^21^ These problems could be overcome by carrying out REHT simulations of protein-membrane-water systems, where we heat the solvent and membrane minimally and heat the protein to a greater extent. In theory, the minimal heating of the membrane should fasten the lateral dynamics of lipids and allow the membrane-bound protein to explore different states. Nonetheless, we must test if REHT can efficiently sample protein binding to the membrane.

Recently, Klimov and co-workers applied REHT on short helical peptide (PGLa) and showed efficient sampling of the peptide binding to anionic bilayer as compared to the REST methods.^15^ Our work, which is similar to the one by Klimov and co-workers, aims to assess the efficiency and robustness of REHT in exploring protein-membrane binding poses as compared against the well-established reference data arrived at by exhaustive AAMD simulations. We also wanted to see if the REHT, which was designed to explore protein conformations, allows one to observe change in protein secondary and tertiary structure as a function of distance from the membrane a feature that would be captured by a larger PMP with some tertiary structure. In order to achieve it, we have chosen the amphipathic lipid-packing sensor (ALPS) motif of Osh4, an oxysterol-binding protein in yeast.^22–24^ The ALPS motif of Osh4 is made up of 27 residues in the N-terminal end of the protein, one of the latter’s six membrane binding regions.^23^ The binding mechanisms of Osh4 ALPS peptide to various membrane compositions were investigated using classical MD simulations by Monje-Galvan and Klauda.^25^ To compare the utility of REHT, we sought to replicate the bound conformations of the peptide with the simplest model membrane used in their study.

The rest of the paper is organized as follows. In the next section following this Introduction section, we briefly discuss the methods and the systems set up. We then discuss the various results showing the performance and accuracy of the method. We finish the paper with a short conclusion.

## 2 Material and Methods

### 2.1 System setup and simulation

The membrane composition used consisted of a mixture of 1,2-dioleoyl-sn-glycero-3-phosphocholine (DOPC) and 1,2-dioleoyl-sn-glycero-3-phospho-L-serine (DOPS) in a 3:2 ratio in bilayers of 80 lipids per leaflet. This mimics the composition of anionic lipids in yeast membranes. The protein-membrane system was created using the Membrane builder module in CHARMM-GUI.^26–28^ Osh4 ALPS motif (27 amino acids) was extracted from the PDB ID: 1zhz^29^ and was horizontally placed at least 6 above the model membrane. Even though Monje-Galvan and Klauda used both vertical and horizontal starting orientations, we have decided to use one of them as the protein would explore different orientations across the replicas in REHT. Water patches of 25 consisting of the TIP3P model^30^ were placed above and below this protein-membrane setup to hydrate the system. The system was neutralized with 64 potassium ions. Energy minimization and equilibration were carried out using GROMACS 2020.4. ^31^ The system was well equilibrated using the CHARMM-GUI six-step protocol^28^ with two NVT steps and four NPT steps for a total time of 15 ns. The protein was restrained throughout this period, and the restraint was slowly removed during this six-step process. The simulation was carried out using the CHARMM36m force field.^32^ The structure at the end of the equilibration, given in Fig. 1(A), was used as the starting structure for the REHT simulation.

**Figure 1:**
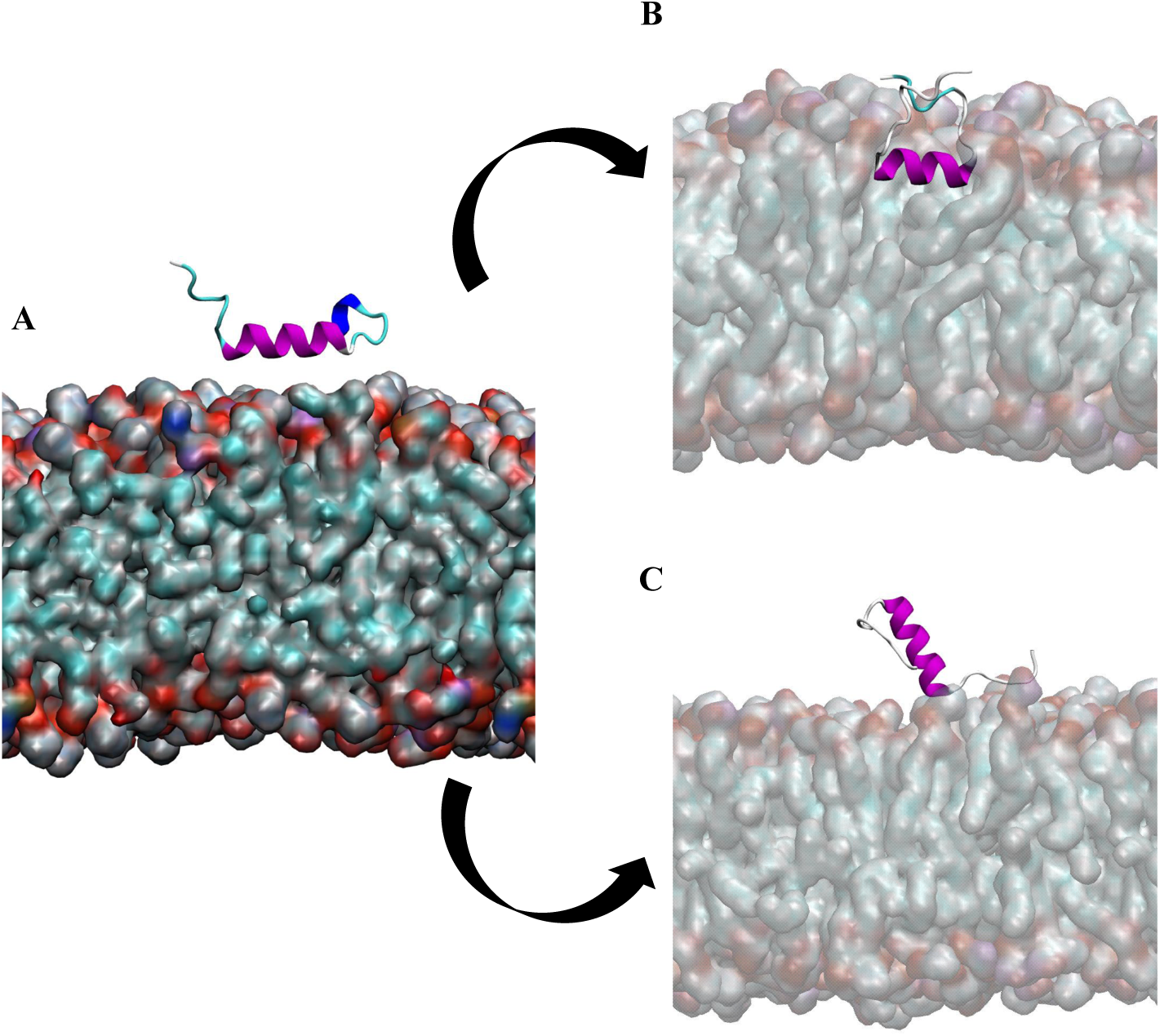
(A) Starting setup for the REHT simulations. The main helix of Osh4 ALPS peptide is marked in purple color. (B) Horizontal and (C) Vertical bound conformations seen in the last 100 ns of the base replica trajectory.

### 2.2 REHT simulation

REHT simulations were carried out with GROMACS 2021.5 patched with PLUMED 2.7.5.^31,32,33^ A total of fifteen replicas were used. The temperature was scaled from 310 to 340 K. This scaling affects all the atoms in the system; hence, the membrane and solvent experience temperatures between 303.15 and 333.15 K across the fifteen replicas. The Hamiltonian of the protein was scaled from 303.15 to 423.15 K. Therefore, the protein experiences both the Hamiltonian and temperature scaling. The simulations used the CHARMM36m force field parameters and the TIP3P water model.^30,32^ The production run was performed in NPT ensemble using a timestep of 2 fs. The hydrogen bonds were constrained using the LINCS algorithm.^35^ Non-bonded forces were calculated with a 12 cutoff (10 : switching distance). Long-range electrostatic forces were calculated using the particle mesh Ewald method.^36^ The temperature of the system in each replica was maintained at their respective temperature and pressure of 1 atm using a Nose–Hoover thermostat^37^ and Parrinello–Rahman^38^ (with semi-isotropic coupling) barostat with time constants 1.0 and 5.0 *ps*^^1^, respectively. Before the exchange in REHT started, all replicas were equilibrated for 1 ns at their respective temperatures, where no exchanges were done. Following this, the production run with replica exchanges was started, and exchanges were attempted every 1 ps. Each replica was simulated for 200 ns, leading to a total simulation time of 3 *µ*s.

## 3 Results and discussion

As mentioned earlier, our main aim of the study was to investigate the utility of REHT in elucidating the membrane-bound conformations of a membrane-binding peptide, Osh4 ALPS peptide. Our findings show that REHT predicts all the membrane-bound conformations in correct populations and captures all the vital residue-wise contacts in those populations. Furthermore, REHT was able to sample these interactions at a much lower simulation time than the classical MD simulations.

### 3.1 REHT performance

We performed the REHT simulation with fifteen replicas, and the exchange probabilities varied between 11-13%. Although this is on the lower side of the accepted range, other analyses given below show enough exploration of all Hamiltonian temperatures (referred to as just ’temperatures’ hereafter) by each replica. The REHT technical performance was analyzed by monitoring the random walk of replicas across Hamiltonian temperatures. The standard demuxing script provided by PLUMED was used to obtain the random walk across temperatures.^34^ Fig. 2(A) shows the random walk of the first and last replicas, and they explore all the temperatures and are not trapped in any temperature for an extended time. Furthermore, Fig. 2(B) shows that all the replicas explore all the temperatures. Although a couple of replicas get trapped in a window of 2-3 temperatures for a few nanoseconds, they come out and explore all the other temperatures during the simulation. To provide a quantitative assessment of the random walk of replicas, the replica mixing parameter was used.^39,40^ It is defined as,

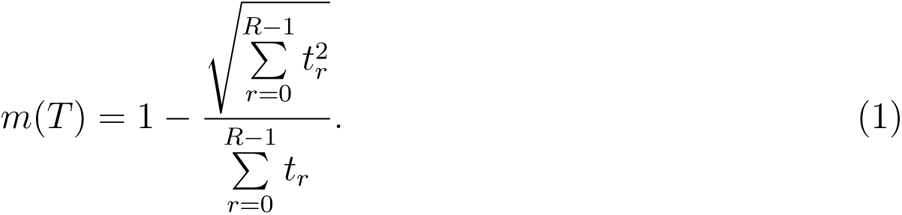

where T is the Hamiltonian temperature, and *t_r_* is the REHT simulation time spent by replica r at temperature T. If we assume an ideal random walk, m(T) reaches a theoretical maximum of 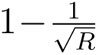, which is constant for all temperatures. ^40^ For our system with 15 replicas, the theoretical maximum is approximately 0.74. Fig. 2(C) shows that the mixing parameter reaches the theoretical maximum for most temperatures except the extreme ones, which could be attributed to the boundary effects. To further establish the adequacy of exchange between temperatures, we plotted the potential energy of each replica, shown in Fig. 2(D). We observe adequate overlap of the potential energy distribution of each replica with its adjacent replicas, which suggests a healthy exchange between them. Having established the good performance of REHT, we characterized the membrane binding properties of the Osh4 ALPS peptide. All the following results were analyzed using the last 100 ns trajectory of the base replica, the replica with a system temperature of 303.15 K in the system temperature scaling.

**Figure 2:**
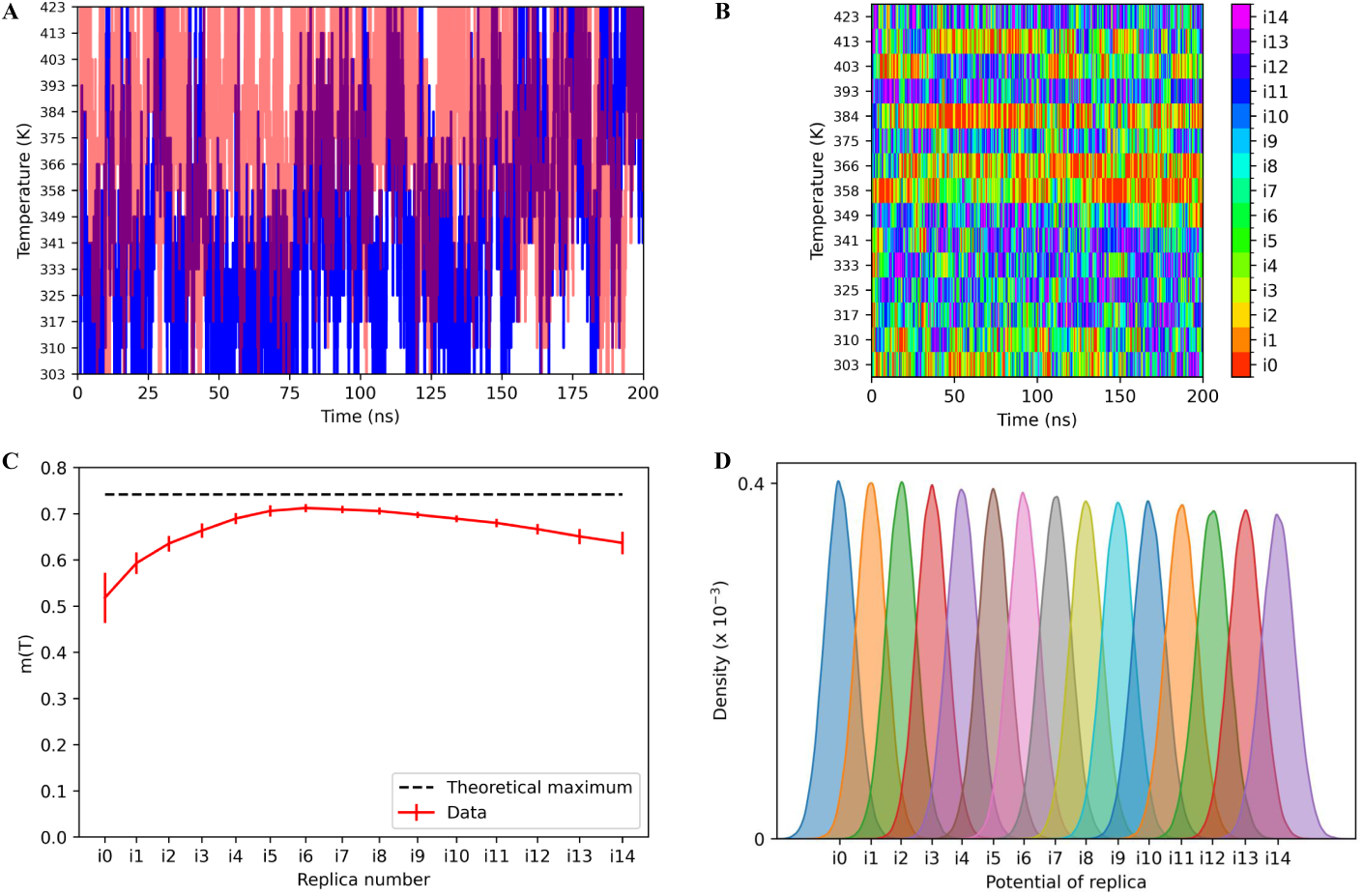
(A) Walk of first and last replicas over Hamiltonian scaling temperature in REHT. First and last replicas are marked in blue and red, respectively. (B) Walk of all replicas over Hamiltonian scaling temperature in REHT. The colors are marked in the color bar on the right. (C) Mixing parameter m(T) averaged over all replicas as a function of Hamiltonian scaling temperature. The theoretical maximum value is marked in black dotted line. Errors are given by red error bars. (D) Overlap of potential energy of replicas. The potential energy distributions of each replica are plotted along the x-axis where the replica numbers are marked.

### 3.2 Osh4 ALPS peptide explores both vertical and horizontal binding poses

First, the Visual Molecular Dynamics (VMD) was used to visually examine the ensemble of structures explored in the base replica trajectory.^41^ Note that it is not a continuous trajectory but an ensemble of states obtained from exchanges across replicas. Qualitatively, we observe that the Osh4 ALPS motif has many membrane-bound states throughout the 200 ns trajectory. However, the membrane-bound peptide was unfolded in many states, which makes up the part of the membrane-bound conformational landscape. Monje-Galvan and Klauda did not report any unfolding of the entire main helix (residues 8 – 18).^25^ For compariosn with their data, we only considered the membrane-bound conformations where the main helix was intact with at least two helical turns. To match this criterion and select only the frames with alpha-helix in the main helix region, we used the Define Secondary Structure of Proteins (DSSP) algorithm.^42^ On examining those selected bound conformations, we notice two different binding modes, horizontal (Fig. 1(B)) and vertical (Fig. 1(C)). To quantify the binding modes, we used the main helix’s tilt angle, defined as the angle subtended by the line joining the C-alpha atoms of residues 8 and 18 with the membrane surface. Ideally, a tilt angle near 0 denotes horizontal binding mode, while an angle near 90 denotes vertical binding mode. However, based on the values reported in Monje-Galvan and Klauda, we decided to use 28 as the cutoff angle for classification into horizontal and vertical binding modes.^25^ Any angle below and above 28 were classified as horizontal and vertical binding modes, respectively. The tilt angle was computed for the selected frames in the last 100 ns of the base replica trajectory. Fig. 3 shows the tilt angle distribution, and we can clearly observe two major regions, one each for horizontal and vertical binding modes. Monje-Galvan and Klauda reported average tilt angle values of 14.73 ± 2.19 and 39.62 ± 4.34 degrees for the horizontal and vertical biding modes.^25^ We also observe peaks for the two binding modes near the reported values in our tilt angle distribution.

**Figure 3:**
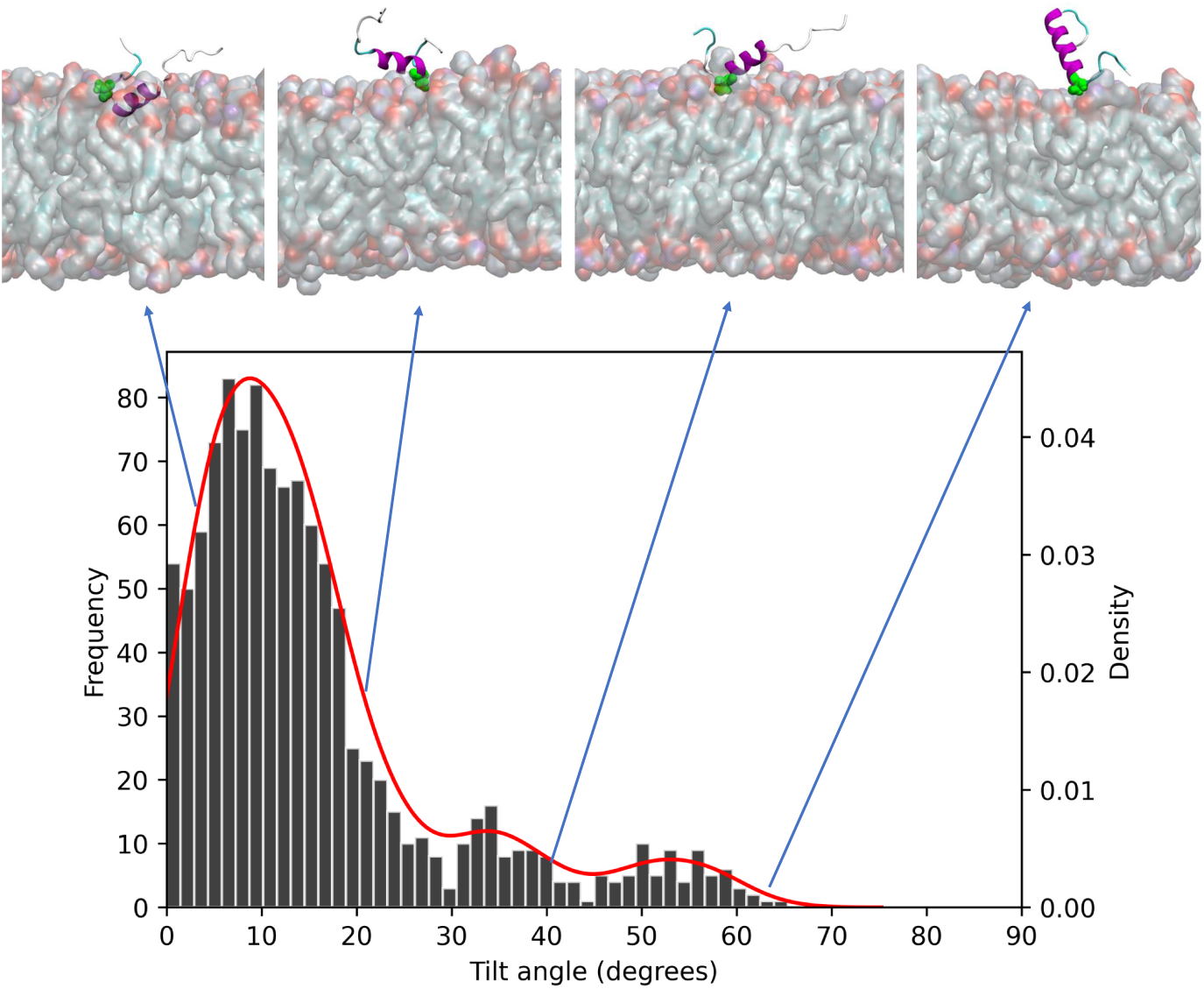
Histogram (Black bins) and density plot (Red line) of tilt angle of membrane-bound helical Osh4 ALPS peptide. The scales for frequency of histogram and density of density plot are marked on the left and right sides of y-axis, respectively. Snapshots of membrane-bound peptide conformations with tilt angles of 5, 22, 40, and 65 degrees are shown above the plot from left to right in order. The main helix of Osh4 ALPS peptide is marked in Purple color and SER8 is marked in Green color.

Furthermore, the population of the horizontal binding mode is much greater than the vertical binding mode. Even though we cannot use the enthalpic component of the binding free energy to validate the population, we can make reserved comments. Monje-Galvan and Klauda have reported average interaction energies of *−*60.46 ± 14.21 and *−*124.17 ± 9.59 kcal/mol for the vertical and horizontal modes, respectively. ^25^ We also observe a similar trend in interaction energies, given in Fig. 4(A). We obtained average interaction energies of *−*86.20 ± 53.09 and *−*198.24 ± 43.35 kcal/mol for the vertical and horizontal modes, respectively, higher than the values reported in classical MD simulations. The higher energies were expected as the base replica visited systems with higher Hamiltonian, increasing the peptide’s interaction energy with the membrane. In essence, both simulations capture the general trend in peptide-membrane interaction energy. Moreover, the skewed population towards the horizontal binding mode that we observe in the tilt angle distribution (Fig. 3) correlates well with such a massive difference in interaction energy. From the populations we obtained in REHT simulations, we calculated the free energy of binding with the tilt angle as the collective variable (Fig. 4(B)), and we see two minima, one each for horizontal and vertical binding modes. The profile we obtained matches the trend in the interaction energies well. However, the difference in free energy between the two binding modes is just 2 kT, and they are further separated by a free energy barrier of 1 kT. Therefore, transitioning from vertical to horizontal binding mode is frequent.

**Figure 4:**
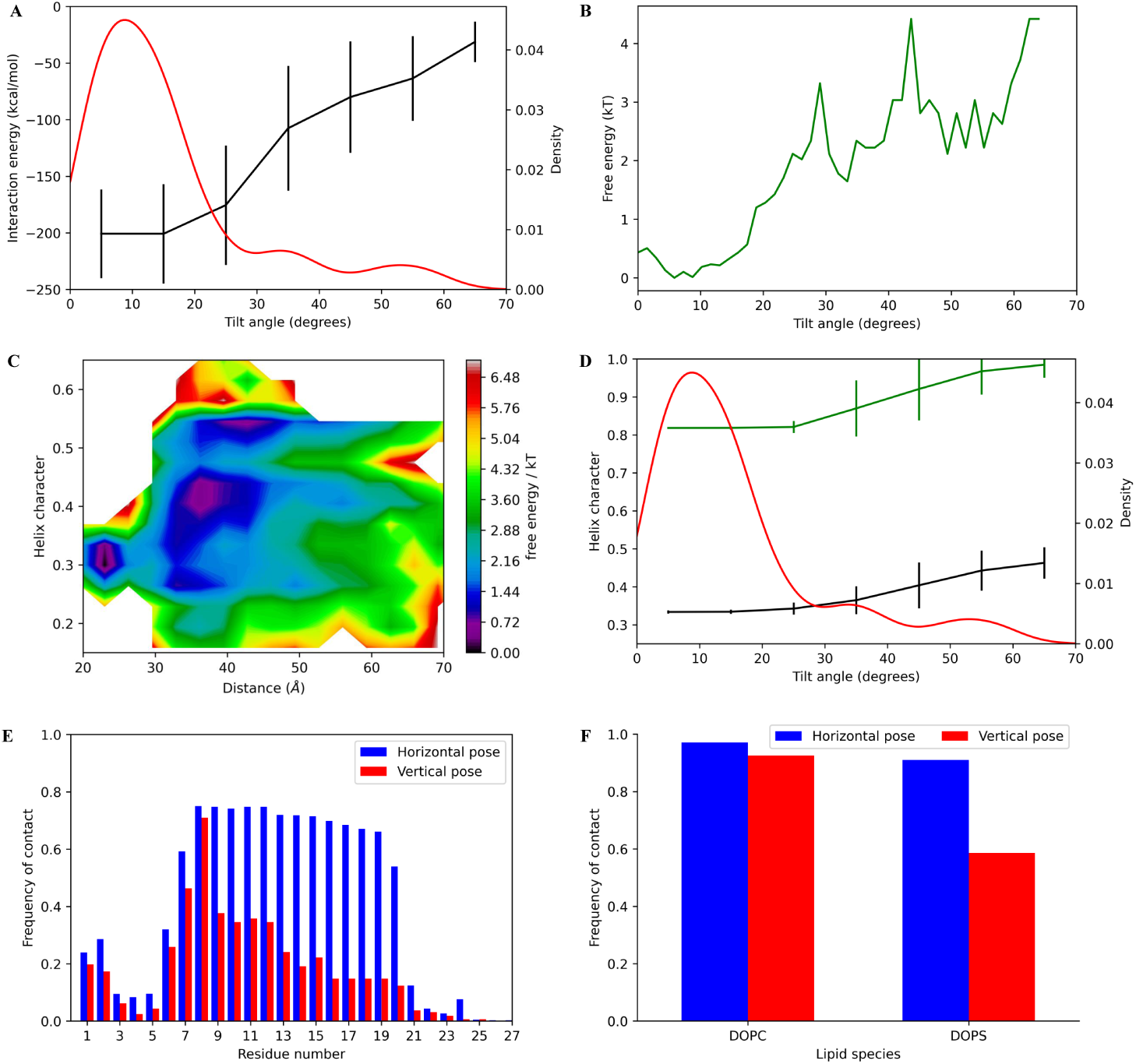
(A) Interaction energy of the Osh4 ALPS peptide with the membrane (black line) as a function of tilt angle. Errors are given by black error bars. The density plot (in red) is given for comparison. (B) The free energy of membrane-bound helical Osh4 ALPS peptide as a function of tilt angle. The free energy is given in kT units. (C) Free energy map of helical character of Osh4 ALPS peptide as a function of the distance of peptide COM from the membrane center. (D) Helical character of the full peptide (black line) and main helix (green line) of the membrane-bound Osh4 ALPS peptide as a function of tilt angle. Errors are given by black and green error bars. The density plot (in red) is given for comparison. (E) Frequency of contact of C-alpha atom of each residue with phosphorous atom in vertical and horizontal binding poses, given in red and blue, respectively. (F) Frequency of contact of the peptide with individual lipid species of the membrane. Vertical and horizontal binding poses are marked in red and blue, respectively.

### 3.3 REHT captures conformation variations as a function of membrane approach

Monje-Galvan and Klauda reported the loss of helicity of the peptide when binding to the membrane with the composition of the yeast membrane. They observed that the peptide partially unfolded before binding and regained helicity on binding, and this happens majorly with the last turn of the main helix.^25^ Although they did not report such partial loss of helicity in the case of the PC-PS membrane, we observed a loss of helicity, especially with the horizontal binding mode. Visually, we observe more helical turns in the vertical binding mode (Fig. 1(C)) than horizontal mode (Fig. 1(B)). To capture the helical character of the peptide as it approached the membrane, we calculated the two-dimensional (2D) free energy profile of the helical character and the center of mass (COM) distance of the peptide from the membrane. Fig. 4(C) shows two major minima, one each for horizontal and vertical binding modes. The minimum nearer the membrane (COM distance of 23) corresponds to the horizontal binding pose and has a lower helicity than the minimum away from the membrane, which corresponds to the vertical binding mode. To differentiate the helicity between the two binding modes, we plotted the helical character of the peptide only for the selected frames with membrane-bound peptide and intact main helix. Again, we clearly observe a difference in the helical character between the two binding modes. Fig. 4(D) shows that the horizontal binding mode has a decreased helical character compared to the vertical binding mode, and this decrease comes mainly from the main helix region (green plot in Fig. 4(D)). As reported by Monje-Galvan and Klauda, we observed the loss of the last turn of the main helix. However, we do not observe any minimum where there is higher helicity with horizontal mode, which implies the lack of refolding seen by Monje-Galvan and Klauda. ^25^

### 3.4 REHT captures the contact frequency between peptide and lipids

To quantify the Osh4 ALPS peptide binding to the membrane, we performed the frequency of contact (FOC) analysis. A contact was said to be present if the C-alpha atom of any residue reached or went below the phosphate plane of the membrane. We observed that the horizontal binding mode had more and longer contacts (Fig. 4(E)). However, the FOC we observed was higher than the values reported by Monje-Galvan and Klauda. ^25^ We believe this disparity in the fractions could be due to the difference in sampling techniques. Nonetheless, our simulations capture all the vital contacts. In both the binding modes, we correctly captured the importance of the SER 6-9 region and the residues at the N-terminal end. Although Monje-Galvan and Klauda reported barely any contact in the region TRP10-LYS15, we observed significant contacts in this region, especially for the horizontal mode. This emphasizes the importance of the contacts made by THR11 and LYS15, as has already been reported. We also calculated the FOC of the peptide for each lipid species. Consistent with the reported results, we find that the horizontal binding mode establishes more number of contact with both the lipid species than the vertical mode (Fig. 4(F)). In the horizontal binding mode, both DOPC and DOPS interacted with the peptide equally, emphasizing the role of electrostatic interactions in the binding process. The same was not observed in the vertical mode, where DOPC had more contacts than DOPS. In comparison, Monje-Galvan and Klauda reported equal binding to both lipid species in both binding modes.^25^

### 3.5 REHT is efficient in sampling the membrane-bound protein conformations

Monje-Galvan and Klauda performed six simulations with DOPC:DOPS membrane (NPT ensemble), amounting to 12 *µ*s of total simulation time. ^25^ On the other hand, we carried out REHT simulations of 200 ns for each of the 15 replicas, leading to a total simulation time of just 3 *µ*s. REHT has the advantage that the binding poses are not user-defined and the algorithm explore the different conformation of binding indepently and exhaustively. In such a short simulation time of 3 *µ*s, we clearly captured both the binding poses reported earlier. Furthermore, we could capture them in a single simulation trajectory (of the base replica) instead of using multiple replicates that gave the range of binding poses with Boltzmann’s weighted populations. This is an essential advantage of replica exchange as we could overcome significant energy barriers that prohibit shifting from one binding pose to another. Another advantage of REHT is the rapid phase space exploration, which leads to quick contact with the membrane. The peptide took just ∼ 25 ns to make contact with the membrane in REHT simulations compared to ∼ 300 ns in classical MD simulations.^25^

## 4 Conclusion

We tested the utility of the REHT method in exploring the equilibrium properties of the membrane-bound PMP conformations and compared it to the results from AAMD.^25^ To do so, we used the Osh4 ALPS motif as the model system and elucidated its membrane-binding characteristics to a DOPC/DOPS model membrane. We demonstrated the two major membrane-bound conformations of the peptide, namely horizontal and vertical binding modes, four times faster than AAMD simulations. In addition to that, the first contact between the peptide and the membrane is accelerated in REHT. REHT not only sampled the conformations efficiently but also captured the protein-membrane interactions accurately. The free energy profile obtained from the populations observed in the REHT base replica showed that the horizontal conformation was favored. Furthermore, the transition from vertical to horizontal binding mode involved the unfolding of the last turn of the main helix. Although Monje-Galvan and Klauda observed such unfolding only with the yeast membrane, we showed that it happens even with the simple anionic model membrane. Our studies captured the importance of SER6-SER9 residues in initiating the membrane binding for both conformations. All the main helix residues, especially LYS15, are essential to a stable horizontal bound conformation. Not only that, REHT was also able to show the importance of contacts with DOPS lipids in the horizontal conformation. Therefore, we can conclude that REHT is efficient and accurate in characterizing the membrane-binding characteristics of membrane-binding peptides. However, we must test the method for larger PMPs. We note that such a study must involve careful choice of the REHT parameters to capture the membrane binding phenomenon accurately (please see the user-manual in the supporting information file). The parameters must be chosen such that the uncharacteristic unfolding of protein or non-physiological membrane properties does not dominate the sampling process. Even though REHT can sample various protein-membrane landscapes, we must be cautious with their interpretation and choose relevant, physiological, and structural regions.

## 5 Acknowledgement

We thank Dr. Rajeswari Appadura (Indian Institute of Science and Education, Tirupati) for the active discussion related to this project. CN thanks the Ministry of Education, Government of India, for the Prime Minister Research Fellowship (PMRF ID: 0201905) and also thanks Indo-French Centre for the Promotion of Advanced Research (IFC-PAR/CEFIPRA) for the Raman-Charpak fellowship (IFC/ 4152/RCF 2023). AS acknowledges the financial support from the Indian Institute of Science-Bangalore and the high-performance computing (HPC) facility “Beagle” setup from grants by a partnership between the Department of Biotechnology of India and the Indian Institute of Science (IISc-DBT partnership programme). AS also thanks the DST for the National Supercomputing Mission grants (DST/NSM/R&D-HPC-Applications/2021/03.10 and DST/NSM/R&D-HPC-Applications/Extension Grant/ 2023/27) for the HPC support. FIST program sponsored by the Department of Science and Technology and UGC, Centre for Advanced Studies and Ministry of Human Resource Development, India. AS would also like to thank the Teams Science Grant from the DBT-Wellcome Trust India Alliance (Grant number: IA/TSG/21/1/600245) and the DBT National Network Project (NNP) grant (BT/PR40323/BTIS/137/78/2023 grants. This work was initiated through the Matrics grants (MTR/2023/001040) from the Science and Engineering Board (SERB), India and both authors thanks SERB for this support.

## Author contributions

AS conceived the idea. CN and AS designed the research. CN performed the research and analyzed the data. AS supervised the study. CN prepared the first draft of the paper and CN and AS polished it together.

## 7 Data availability statement

All input files related to the REHT simulations of OSH4 systems are curated and publicly available at our laboratory Github repository: codesrivastavalab/REHT-PMP. We have also included a supporting information file as a USER MANUAL for the method.

## Supporting Information

### REHT USER manual

In this manual, we will set up and run ‘Replica exchange with hybrid tempering’ simulations using an already equilibrated system. The manual is written in a generic manner to set up simulations for protein-water system as well as for protein-membrane-water system. All the steps described hold for both types of systems. In this manual, we will use the GROMACS MD engine patched with PLUMED.

All the required files for protein-water system are uploaded at: https://github.com/codesrivastavalab/ReplicaExchangeWithHybridTempering

Input files needed to run peripheral membrane protein system (OSH4) using REHT is available at: https://github.com/codesrivastavalab/REHT-PMP

#### 1. System

We need the following files to begin with:

i. Structure (.gro) file of the equilibrated system {input.gro}
ii. Topology file for your system {topol.top}
iii. Index file for your system with various groups: protein, membrane (if present), and solvent {index.ndx}

##### 2. MDP file generation (Temperature scaling)

In this step, we will generate an ‘N’ number of mdp files, one for each replica. The only difference between the mdp files will be the system’s temperature. Therefore, all the atoms in the system will experience this temperature. It could be solvent or solvent and membrane, depending on your system. Note that the protein also experiences this temperature range in addition to the Hamiltonian scaling (discussed below). The required files are in the “*3_REHT_input_generation*” section of the GitHub page mentioned above.

i. Prepare your mdp file with ‘XXX’ marked in place of the temperature as shown below. The groups depend on our system. All the other inputs can be as per our requirements.

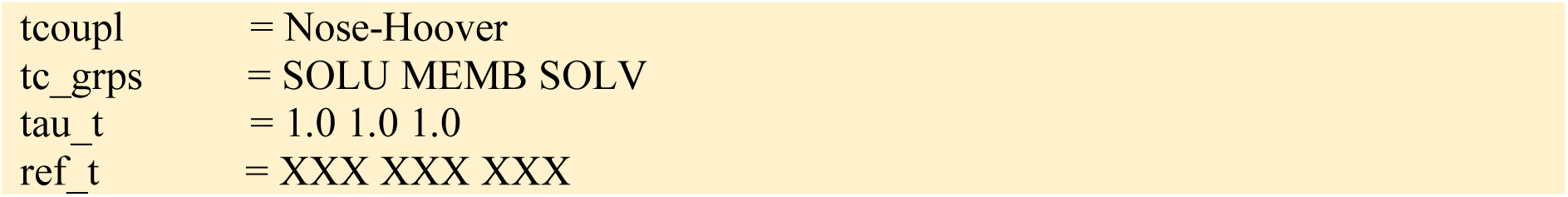
ii. Download and use the “*temp.sh*” file in the above-mentioned section. We must change the number of replicas and the temperature range per our requirements. In this step, we build a set of temperature values using geometric progression in the temperature range of choice. Then, each value is written out in a separate mdp file. For instance, if we use 20 replicas with temperature scaling of 310 K to 340 K, we will obtain 20 mdp files (md0.mdp, md1.mdp, …, md19.mdp) with geometrically progressing temperature in successive replicas.

##### 3. Topology file generation (Hamiltonian scaling)

In this step, we will generate an ‘N’ number of topology files, one for each replica. Here, we will scale the Hamiltonian of the protein alone based on the required temperature range (for example, 310 K to 450 K) for Hamiltonian scaling. Therefore, only the protein experiences this temperature range and not any other atom in the system. We use the “plumed partial_tempering” command in PLUMED to do this.

i. The first step is to obtain a topology file without any include statement (as seen in top files generated from CHARMM-GUI). We use the following command to get that processed topology file: *gmx grompp -f md0.mdp -c input.gro -n index.ndx -p topol.top -pp processed.top*
ii. Once we get the processed.top file, we must mark the “hot” atoms with an underscore, “_”, after the atom type as given below. (“Hot” atoms are the protein atoms that will be subjected to the Hamiltonian scaling). 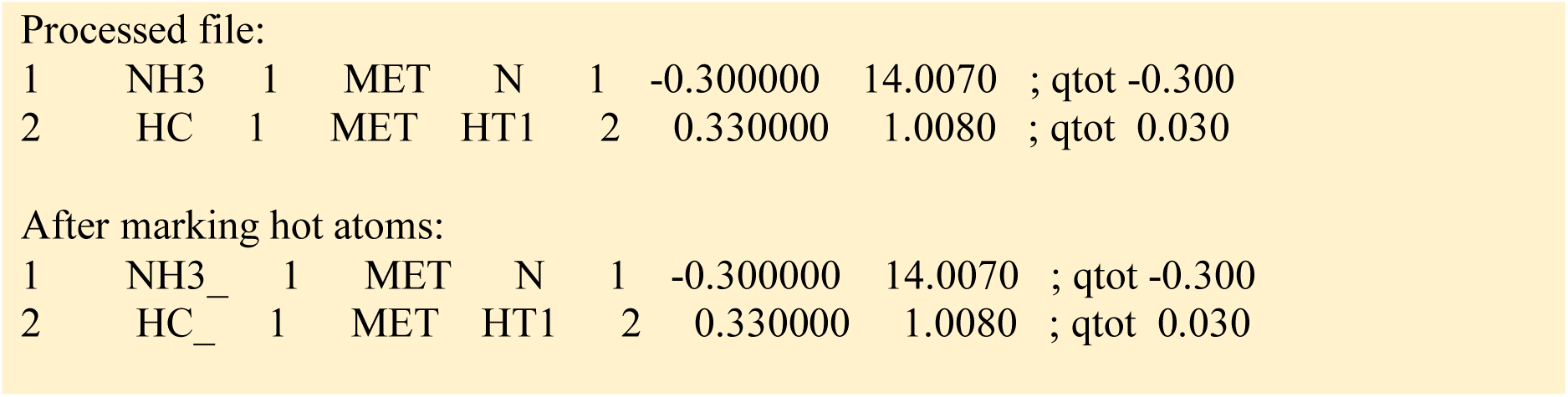 To do so, we should check the processed.top file and note down the starting and ending lines of the protein atoms. Then, we use the following command to generate a new topology file with marked hot atoms: *sed ‘startline,endline s/./_/18’ processed.top > processed_.top*
iii. Let us assume the starting and ending lines are 2016 and 2954, respectively. Then the command would be:
iv. *sed ‘2016,2954 s/./_/18’ processed.top > processed_.top*
v. We then generate the scaled topology files from the ‘processed_.top’ file using the “*scale_topol.sh*” file in the above-mentioned section. Again, we build a set of temperature values using the geometric progression in the temperature range of choice and then use the “*plumed partial_tempering*” command to generate the scaled topology files.
vi. In this file, we have to edit the number of replicas and the temperature range for the Hamiltonian scaling of the protein. We should use one of the two sections depending on the forcefield used.
  a. If ‘charmm36m’ or a forcefield with cmap terms are used, we must comment out line 29 and use the section from lines 32 to 45.
  b. If other forcefields are used, only line 29 is sufficient.

Once we execute this file, we will obtain the ‘N’ number of topology files. If we give 20 replicas, we will obtain 20 files named md0.top, md1.top, …, md19.top.

##### Note on software

The Gromacs 2018 version and before had a “*multi*” option for their “*gmx mdrun*” command. However, this option is no longer available. It has been replaced with the “*multidir*” option in the newer versions. It requires the presence of a separate folder for each replica. It is better to use the latest version with the “*multidir*” option.

#### 4. Equilibration

Now, we have generated the required number of mdp and topology files. We also need a “*plumed.dat*” file, which is usually empty for REHT simulation. However, it can be edited to carry out other advanced sampling techniques.

In our REHT run folder, we must have the following files:

i. Structure file {input.gro}
ii. Index file {index.ndx}
iii. mdp files {md0.mdp, md1.mdp, …, md19.mdp}
iv. Topology files {md0.top, md1.top, …, md19.top}
v. plumed.dat (empty file)

Once we have these files, we can start the equilibration step. In this step, we run each replica for 1 ns to equilibrate the replica to their respective temperature. During this period, we do not attempt any exchange between the replicas.

We use the following script to generate the required number of folders and start the equilibration:

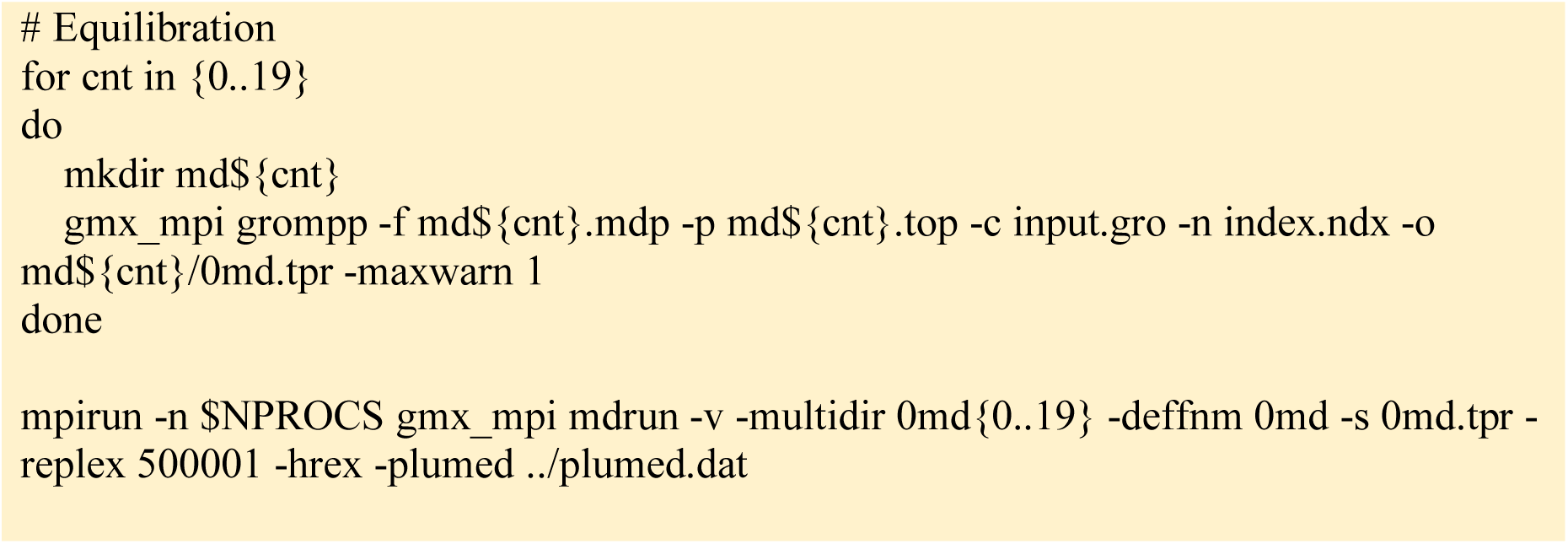

In the “*gmx mdrun*” command, the “*hrex*” option enables Hamiltonian replica exchange, and the “*replex*” option gives the frequency in terms of the number of steps. Since we have given replex as 500001, there should be no exchange in the 1 ns equilibration run.

#### 5. Main run with exchanges

Once we have equilibrated the replicas, we can start the main REHT run with exchanges attempted at a pre-determined frequency. We usually attempt an exchange every 1 or 2 ps. We use the following script to extend the simulation:

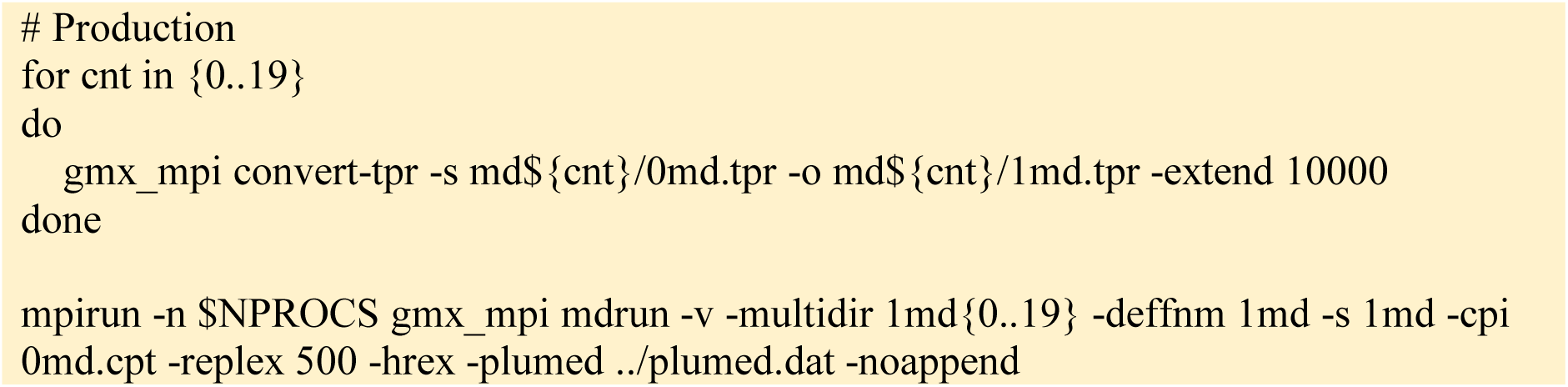

First, we run the REHT simulation for 10 ns to determine if our REHT setup (scaling temperatures and number of replicas) leads to a healthy exchange of about 10-30% across all the replicas. This information (shown below) is given in the log file towards the end.

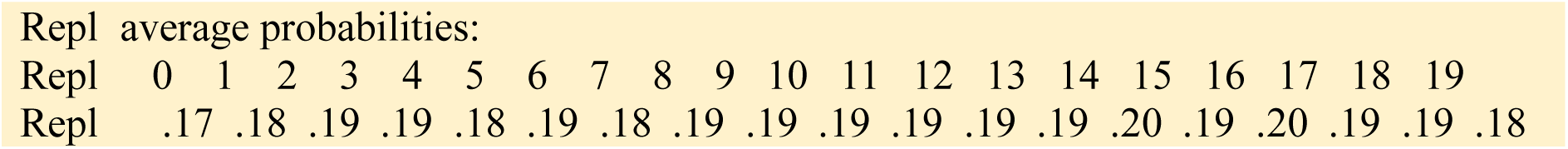

Furthermore, we should also check the random walk of the replicas by “demuxing” the trajectories. To do this, please follow the procedure in the “PLUMED Masterclass 21.5: Simulations with multiple replicas” tutorial page on the PLUMED website. They have explained the procedure and have also uploaded the Perl script to do it. Using it, we should check if all the replicas explore the full range of indices. Sometimes, this exploration may not occur within the first 10 ns and might take about 100 ns.

Once we ascertain our setup, we can extend the simulation to the required time. HAPPY SIMULATING!

